# High-Level Expression, Purification and Biophysical Characterization of GPI-anchored native-like human Prion Protein using *Leishmania tarentolae*

**DOI:** 10.1101/2025.03.30.646161

**Authors:** Najoua Bolakhrif, Thomas Pauly, Luitgard Nagel, Dieter Willbold, Lothar Gremer

## Abstract

The human prion protein (PrP) is a glycosylphosphatidylinositol (GPI)-linked membrane-bound glycoprotein, containing two glycosylation sites. Human PrP is associated with a number of neurodegenerative diseases, called transmissible spongiform encephalopathies (TSE). Pathogenesis involves a structural conversion of the cellular form (PrP^C^), rich in α-helical and random coil structure, into the scrapie form (PrP^Sc^) characterized by parallel in register intermolecular β-sheet conformation. To get a better understanding of this structural conversion, it is crucial to first characterize the non-pathogenic cellular isoform including all posttranslational modifications, like GPI-anchoring and native-like human glycosylation pattern. So far, studies on PrP^C^ or PrP^Sc^ as well as the transition from one state to the other rely on non-native constructs of PrP studied far away from physiological conditions. We, therefore, established the expression of GPI-linked human PrP with close to native glycosylation pattern (native-like human PrP) using the eukaryotic LEXSY expression system in *Leishmania tarentolae*. This expression system has the added advantage that it allows for large-scale production of the native-like human PrP, which results in ∼1 mg purified protein per liter culture. Sedimentation velocity analysis and far-UV circular dichroism spectroscopy confirm the high structural homogeneity and monomeric native-like conformation of the purified GPI-anchored human PrP.

## Introduction

The human prion protein (huPrP) consists of 253 amino acid residues and is a membrane-bound glycoprotein mainly located in the nervous system and containing an unstructured N-terminal domain and a globular C-terminal domain, consisting of mainly alpha-helices. It is most known for its involvement in a number of neurodegenerative diseases including Creutzfeldt-Jakob disease (CJD). These prion diseases are transmissible spongiform encephalopathies (TSEs) involving the abnormal accumulation of misfolded huPrP as amyloid fibrils (Wang et al., 2020). “Prions” are misfolded aggregates of the prion protein that are partially resistant to proteinase K digestion and further have the ability to transfer structural identity to recruit native protein by autocatalytic processes (Prusiner, 1982; Stöhr et al., 2008; Willbold, Strodel, Schröder, Hoyer, & Heise, 2021). The native, or cellular form of huPrP (huPrP^C^) is characterized mainly by α-helical and random coil structural elements (Zahn et al., 2000). The first 22 amino acids of the N-terminal domain are responsible for the transport to the endoplasmic reticulum (ER) (Kim, Rahbar, & Hegde, 2001) and are subsequently cleaved off from the huPrP precursor. Adjacent to the transport signal sequence is an unstructured (random coil) region known as the hydrophobic N-terminal domain which comprises five octameric repeats (amino acids 23 to 127). A small anti-parallel β-sheet (amino acids 128 to 131 and amino acids 161 to 164) represents about 3 % of the total structure (Pan et al., 1993; Zahn et al., 2000). The C-terminal domain has approximately 42 % α-helical structure, including helix 1 (amino acids 144 to 154), helix 2 (amino acids 173 to 194), and helix 3 (amino acids 200 to 228). Lastly, membrane attachment is achieved *via* a glycosylphosphatidylinositol (GPI)-anchor present at the C-terminal end of Ser230 with a molecular weight of approximately 2 kDa (Stahl et al., 1992). GPI anchoring is achieved through the C-terminal GPI anchoring signal sequence comprising amino acids 231 to 253. This signal sequence is afterwards cleaved off from the huPrP precursor. The total size of huPrP varies between approximately 30 and 35 kDa, depending on the number of glycans at glycosylation sites N181 and N197 (excluding the GPI anchor). The native structure of mainly α-helices and random coil holds for the cellular isoform (huPrP^C^), however, the misfolded form (huPrP^Sc^), is dominated by a parallel in register intermolecular β-sheet conformation (Groveman et al., 2014; Hoyt et al., 2022; Kraus et al., 2021; Pan et al., 1993; Safar, Roller, Gajdusek, & Gibbs, 1993). The details of this structural conversion of PrP^C^ into PrP^Sc^ still remain elusive. Sudies suggest that conversion occurs in close proximity to the cell membrane (Vey et al., 1996). Interestingly, experiments on full-length PrP where the GPI anchor was excluded do not show any spontaneous conversion under physiological conditions (Kaneko et al., 1997; Resenberger et al., 2011; Taraboulos et al., 1995). As a consequence, studies focusing on the full-length huPrP utilized destabilization of the native folding by acidic pH or denaturants to induce refolding into the amyloid conformation (Bocharova, Breydo, Parfenov, Salnikov, & Baskakov, 2005; Pauly et al., 2022). An alternative approach has been to exclude the globular C-terminal domain and use shorter constructs of huPrP to facilitate the spontaneous conversion of the unstructured N-terminal region from an unstructured monomeric state into an amyloid state (Kundu et al., 2003). Furthermore, investigations on the conversion of huPrP^C^ into huPrP^Sc^ usually rely on bacterially expressed recombinant huPrP^C^ lacking posttranslational modifications (PTMs). Expression systems involving PTMs commonly utilize *Pichia pastoris*, but their glycosylation patterns diverge from mammalian patterns (Mehlhorn et al., 1996; Riley et al., 2002). Indeed, a successful expression in *P. pastoris* of GPI-anchored PrP from Syrian golden hamster is reported (Marbach, Zentis, Ellinger, Müller, & Birkmann, 2013). To overcome these shortages, we established the expression of the membrane-bound full-length huPrP^C^ with human-like PTMs in the eukaryotic trypanosomatid parasite *Leishmania tarentolae. L. tarentolae* is one of the very few expression systems providing glycosylation patterns resembling those in humans, except for missing the terminal sialic acid (Lai, Klatt, & Lim, 2019; Niimi, 2012). We found *L. tarentolae* to be an appropriate eukaryotic expression system for producing large quantities of homogeneous, fully processed huPrP^C^ for accurate studies on the structure and conversion closely linked to human prion diseases. Several studies support the capability of *L. tarentolae* to yield biologically active eukaryotic proteins as well as mammalian-like N-glycans (Breitling et al., 2002; La Flamme, Buckner, Swindle, Ajioka, & Van Voorhis, 1995; Niimi, 2012; Phan, Sugino, & Niimi, 2009; Zhang, Charest, & Matlashewski, 1995). We introduce here this new expression system for high quality native-like huPrP^C^, including its purification and confirmation of homogeneity by sedimentation velocity analysis by analytical ultracentrifugation (AUC) and far-UV circular dichroism (CD) spectroscopy. This work intends to improve the understanding of PrP misfolding by bringing *in vitro* studies closer to physiological conditions.

## Results

### Vector design and expression of native-like huPrP^C^

The full-length huPrP sequence including the N-terminal and C-terminal signal sequences was cloned into the *L. tarentolae* expression vector pLEXSY-neo2.1. The flexible N-terminal region contains a signal sequence, responsible for the transport to the ER. After the N-terminal signal sequence, we inserted a 6×His tag followed by a factor Xa cleavage site (Figure 1A). This allows for affinity chromatography purification and subsequent cleavage of the 6×His-tag. To guarantee the permanent genomic integration of the huPrP sequence, the expression cassette was integrated into the *odc* locus of chromosome 12 of *L. tarentolae*. The integration was confirmed by diagnostic PCR, after the preparation of genomic DNA from a dense culture, resulting in 1.1 kbp and 2.4 kbp characteristic fragments for the positive clones. Clones that were not genome-integrated did not show bands with respective sizes (Figure 1B). After induction of expression of native-like huPrP with tetracycline, SDS-PAGE and western blot of the cells as well as of the culture supernatant was performed. This revealed intracellular expression with no secretion of the protein into the culturing medium (Figure 1C). Furthermore, a background expression was detected, without induction, enabling the selection by geneticin, G-418. Enzymatic digestion using phosphatidylinositol phospholipase C (PI-PLC), which is a phosphatidylinositol-specific enzyme, was used to confirm membrane anchoring by cleaving off the GPI anchor. This resulted in native-like huPrP detected in the supernatant after centrifugation, while no treatment showed only signal in the cell pellet (Figure 1D).

**Figure 1.**
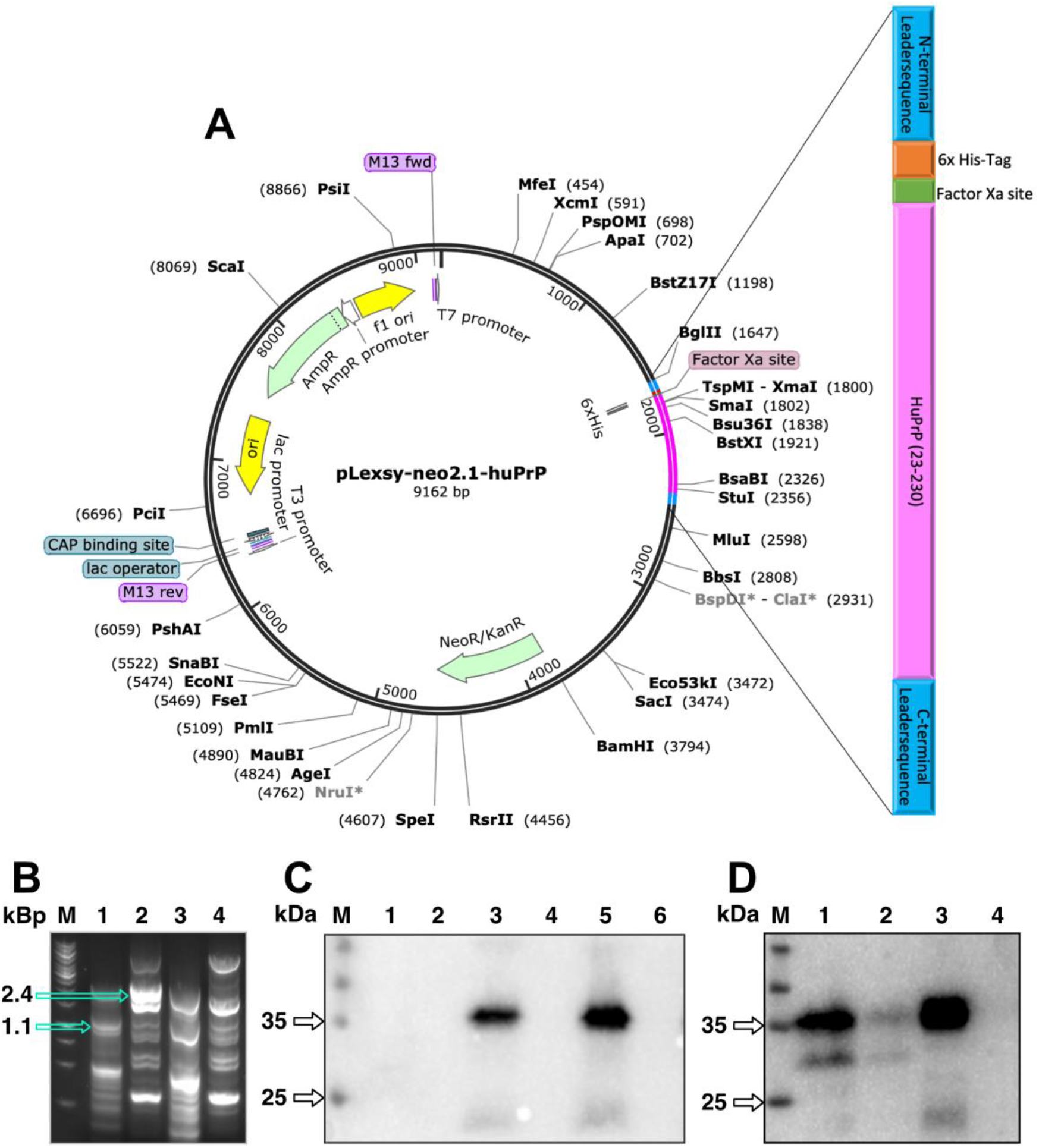
Native-like huPrP: Cloning and expression in *Leishmania tarentolae*. (A) Gene map of the pLexsy neo.2.1 vector including the huPrP sequence, inserted using the restriction sites NcoI/ NotI. (B) Confirmation of genome integration by diagnostic PCR. 1: Positive clone, control region: 5’ odc – utr1 resulting in fragment size of 1.1 kBp; 2: Positive clone, control region: neo – 3’odc resulting in fragment size of 2.4 kBp; 3: Negative clone, control region: 5’ odc – utr1; 4: Negative clone, control region: neo – 3’odc. (C) Evaluation of huPrP protein expression by western blot. 1: cellular detection of huPrP in *L. tarentolae* before vector integration; 2: secretory detection of huPrP in *L. tarentolae* before vector integration; 3: cellular detection of huPrP in *L. tarentolae* after vector integration and before induction with tetracycline; 4: secretory detection of huPrP in *L. tarentolae* after vector integration and before induction with tetracycline; 5: cellular detection of huPrP in *L. tarentolae* after vector integration and after induction with tetracycline; 6: secretory detection of huPrP in *L. tarentolae* after vector integration and after induction with tetracycline. (D) Characterization of post-translational modifications of huPrP expressed in *L. tarentolae* by western blot. 1: Pellet of *L. tarentolae* expressing native-like huPrP after 1 h treatment with PI-PLC; 2: Supernatant of *L. tarentolae* after 1 h treatment with PI-PLC; 3: Pellet of *L. tarentolae* expressing native-like huPrP without treatment with PI-PLC; 4: Supernatant of *L. tarentolae* without treatment with PI-PLC.

Moreover, the western blot revealed multiple bands between 25 kDa and 35 kDa, which are characteristic for cellular and glycosylated huPrP (Figure 1D). Lastly, proteolysis of the expressed native-like huPrP with proteinase K shows no resistance to digestion supporting the cellular conformation (Suppl. Figure 1).

### Solubilization and purification of native-like huPrP

Since huPrP is linked to the membrane *via* a GPI anchor, differential centrifugation was applied after cell disruption to obtain the membrane fraction. A final step of 100,000 x*g* is applied to pellet the membrane fraction (see methods for further details). To prevent aggregation, membrane proteins are usually transferred from the native cellular membrane to membrane mimetics. Several detergents were tested for the ability to solubilize native-like huPrP (Suppl. Figure 2). n-Dodecyl-β-D-maltoside (DDM) is the detergent of choice. It is a mild and non-denaturing detergent, which is often able to preserve the native protein folding, with a relatively high critical micellar concentration (e.g. 0.12 mM in 0.2 M NaCl), which is crucial for subsequent removal and replacement of the detergent. Lastly, the optical properties of DDM do not interfere with most spectroscopic methods commonly used to study protein structure and hydrodynamic properties, as in the case for Triton X-100 or NP-40. After solubilization, native-like huPrP was purified using immobilized metal affinity chromatography (IMAC).

Applying an imidazole gradient for elution resulted in a prominent peak containing native-like huPrP (Figure 2A). The fractions corresponding to the peak from IMAC were then subjected to size exclusion chromatography (SEC) where two major peaks with retention volumes of approximately 10 ml and 15 ml on Superdex 200 Increase 10/300 were seen (Figure 2B). Using a dot blot, the second peak was found to contain native-like huPrP (Suppl. Figure 3). The purity after each step was analyzed by semi-denaturing SDS-PAGE and subsequent Coomassie staining. After IMAC, a prominent band at around 40 kDa, corresponding to huPrP, is present (Figure 2C). After SEC, most of the remaining impurities were removed resulting in only one apparent band at approximately 40 kDa. It should be noted that since huPrP is incorporated via its GPI anchor in DDM micelles, the samples could not be heated to 95 °C, as is commonly done in preparing samples for SDS-PAGE but were instead loaded onto the gel directly after the addition of reducing SDS-PAGE sample buffer. The consequence is a slightly increased molecular mass of huPrP (40 kDa versus 30-35 kDa), which is a typical hallmark for micelle-bound membrane proteins (Rath, Glibowicka, Nadeau, Chen, & Deber, 2009) (Figure 2C). UV absorbance spectroscopy of the purified native-like huPrP shows a distinctive profile with two local maxima at 273 nm and 283 nm, while the full-length huPrP expressed in *E. coli* shows a characteristic profile with one major peak at 280 nm (Suppl. Figure 4). We ascribe the UV absorbance differences of purified native-like huPrP *vs*. full-length huPrP expressed in *E. coli* due to its presence (in native-like) or absence of the GPI-anchor (expressed in *E. coli*). The described purification protocol yielded 0.63 mg of highly pure native-like huPrP from a 600 ml *L. tarentolae* culture. The hydrodynamic properties and protein conformation were further analyzed using biophysical methods.

**Figure 2.**
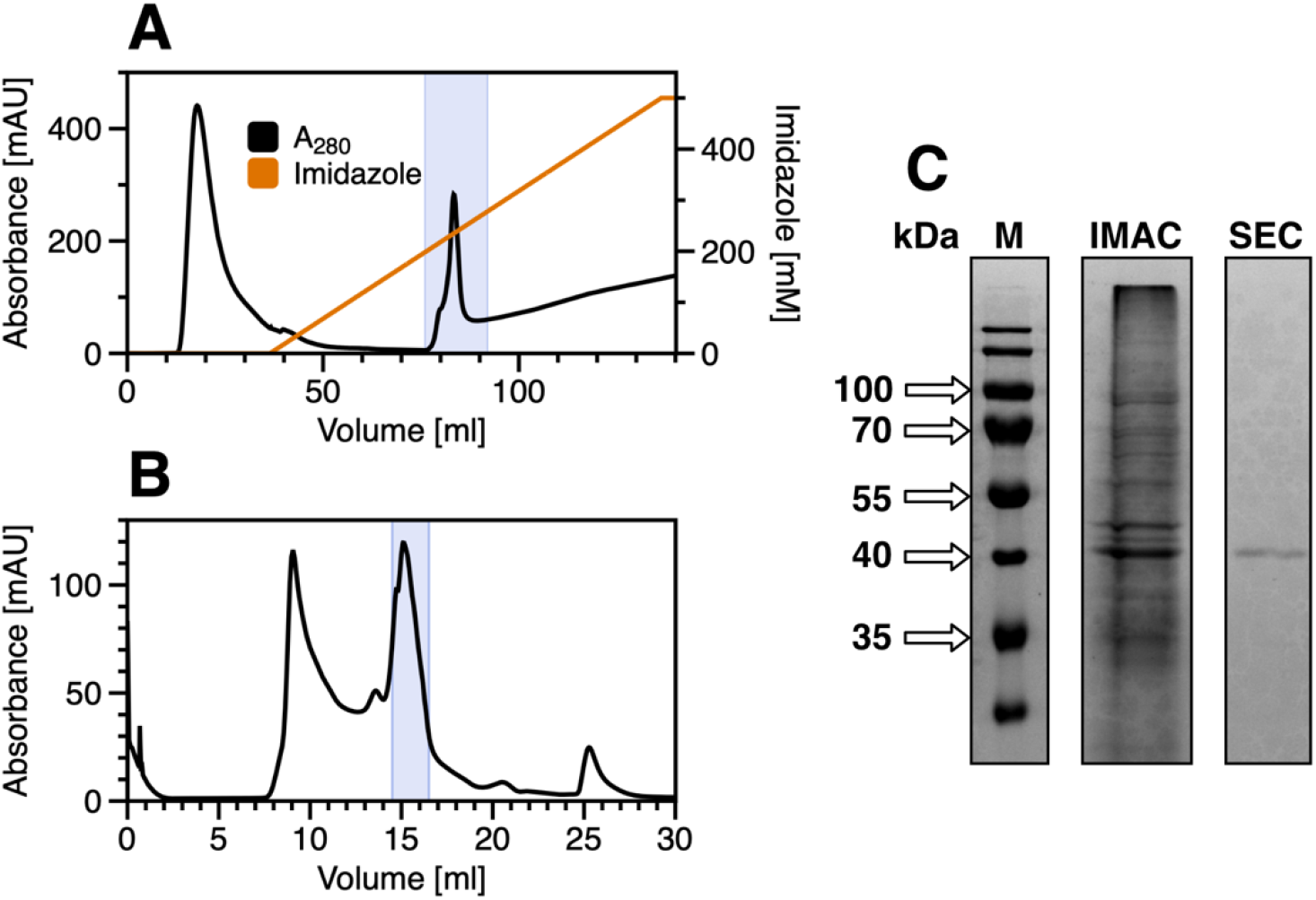
Isolation and purification of recombinant native-like huPrP produced in *L. tarentolae*. (A) Purification by immobilized metal affinity chromatography (IMAC) on a Protino Ni-NTA column after solubilization of the membrane fraction. Absorbance at 280 nm and imidazole concentration is plotted against the elution volume. The peak including huPrP is highlighted in light blue. (B) Purification of the huPrP containing fraction after IMAC by size exclusion chromatography (SEC) on a Superdex 200 Increase (10/300) column. Absorbance at 280 nm is plotted against the elution volume. The peak containing native-like huPrP is highlighted in light blue. (C) Determination of purification by Coomassie-stained SDS-PAGE of huPrP containing fractions (highlighted in A and B in light blue) after IMAC and SEC.

### Biophysical characterization of native-like huPrP

When studying recombinant huPrP from *E. coli*, it is common to reduce the pH considerably away from physiological relevant values, in order to ensure solubility (López García, Zahn, Riek, & Wüthrich, 2000; Riek et al., 1996). It should be noted that the secondary structure of recombinant huPrP from *E. coli* at pH 4.5 was previously studied by NMR spectroscopy, which revealed that the protein mainly adopted a structure composed of α-helical and random coil elements with a single small antiparallel β-sheet (Zahn et al., 2000), in line with proposed structures at native conditions (López García et al., 2000; Riek et al., 1996). To investigate the secondary structure of purified native-like huPrP from *L. tarentolae* solubilized in DDM micelles, far-UV CD spectra were recorded under near-physiological conditions (50 mM Tris-HCl, 150 mM NaCl, pH 7.4 as buffer system, containing 0.02 % DDM (w/v)) and compared to full-length huPrP expressed in *E. coli* under similar conditions (Figure 3A). The far-UV CD spectrum of native-like huPrP shows negative bands at 208 nm and 222 nm, typical for α-helical structure (Johnson, 1988; Kardos et al., 2025). Solvent absorbance at low wavelengths impeded analysis of typical bands below 200 nm. As disordered proteins have a slightly positive signal above 210 nm, this structural element could contribute to the observed far-UV CD spectrum, which reports on weighted average protein structure. In comparison, full-length huPrP from *E. coli* was not fully soluble using similar buffer conditions with physiological ionic strength (50 mM Tris-HCl, 150 mM NaCl, pH 7.4). Therefore, the experimental conditions were adjusted to 10 mM Tris-HCl, pH 7.4 without NaCl for far-UV CD measurements of full-length huPrP from *E. coli*. Full-length huPrP shows a higher fraction of β-sheet structure under these conditions, indicated by a shift of the two negative bands closer to 218 nm, which is the typical minimum for β-sheet containing structures. The structure comprises a mixture of α-helical, random coil and β-sheet structure with an increased β-sheet content compared to native-like huPrP. The hydrodynamic properties of native-like huPrP and full-length huPrP were studied by sedimentation velocity (SV) experiments at near-physiological conditions (50 mM Tris-HCl, 150 mM NaCl, pH 7.4 as buffer system) (Figure 3B, C, and D). The majority (86.4 %) of native-like huPrP is detected at an apparent sedimentation coefficient (*s*-value) of 2.96 S with a calculated mass appropriate for the monomeric protein. The remaining signal accounts for a larger species at 4.94 S with a calculated mass appropriate for a dimer. In contrast to native-like huPrP from *L. tarentolae*, full-length huPrP from *E. coli* suffers from reduced solubility at physiological conditions. Full-length huPrP showed a majority of large aggregates (58.3 % of total signal), which was determined as a loss of signal during the acceleration of the centrifuge. The major part of the remaining soluble fraction (21.8 %) was detected at 1.81 S with a calculated mass appropriate for the monomeric. Therefore, both far-UV CD and SV AUC analyses clearly show that purified native-like huPrP expressed in *L. tarentolae* is superior to full-length huPrP expressed in *E. coli* for studying huPrP in its monomeric native state.

**Figure 3.**
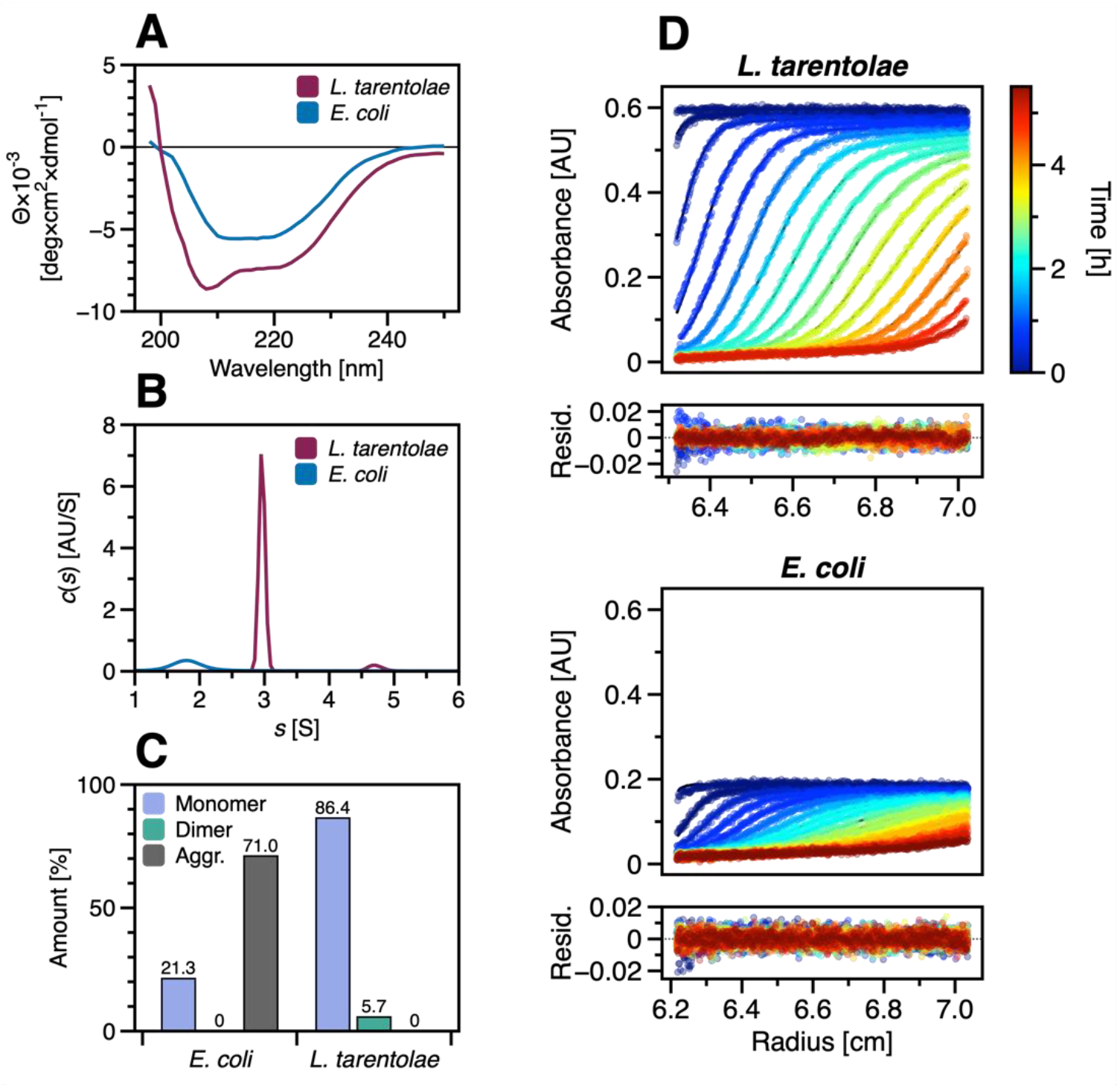
Analytical comparison of huPrP obtained from *E. coli* or *L. tarentolae*. (A) Far-UV CD spectra of 10 μM of huPrP from *L. tarentolae* (purple) and *E. coli* (blue). (B) The result of fitting sedimentation profiles is a distribution of *s*-values. (C) Integration of *s*-value distributions yields the respective monomeric fractions of the sample. The loss of signal during acceleration of the centrifuge gives rise to the aggregate amount. (D) Sedimentation profiles of 10 µM huPrP show raw data as points with fitted Lamm-equation solutions as lines from SV-AUC experiments. The timestamp of each scan is color-coded.

## Discussion

The most frequent prion disease in humans is sporadic CJD, which involves the conversion of huPrP^C^ into huPrP^Sc^ (Zerr & Parchi, 2018). However, genetic predisposition is known to affect susceptibility to certain prion diseases (Palmer MS, Dryden AJ, Hughes JT, & Collinge J, 1991). To understand the onset of the disease, it is crucial to understand the mode of action in detail that leads to the structural conversion of the native protein. The pathological isoform, PrP^Sc^ has been investigated by several studies revealing its highly stable amyloid structure (Artikis, Kraus, & Caughey, 2022; Hoyt et al., 2022; Kraus et al., 2021; Wang et al., 2020). Most of these studies suffer from a trade-off involving two drawbacks: First, high protein yields are available from recombinant production using *E. coli*, which consequently lacks conformity with PrP found in humans; second, relevant PTMs are present, which usually come with high costs or low protein yields, e.g. for CHO cells, 6–10 μg Chinese hamster ovary (CHO)-PrP^C^/g CHO cells (Elfrink & Riesner, 2004). Another approach, which is no less important, focuses on the structural properties of the cellular isoform and its misfolding. This approach faces further obstacles to overcome, including the solubility of full-length huPrP expressed in *E. coli* at physiological pH values. To avoid these problems, most studies focused on the investigation of shorter constructs of PrP (Gerum, Silvers, Wirmer-Bartoschek, & Schwalbe, 2009; Hosszu et al., 2005), PrP from other species (Eghiaian et al., 2007), or lowered pH (Gerum et al., 2009; Pauly et al., 2022). Despite the fundamental insights that such *in vitro* studies revealed regarding protein structure and misfolding mechanisms, there is an obvious discrepancy to *in vivo* conditions. *In vitro* studies applying high-resolution structural methods are in urgent need of closing the gap to physiological conditions to yield significant advances in understanding prion diseases. So far, NMR studies revealed a predominantly α-helical structure of the cellular isoform, although these experiments were also conducted at an acidic pH (pH 4.5). At physiological pH conditions using far-UV CD spectroscopy, we were unable to confirm the high alpha-helical structure content for full-length huPrP from *E. coli*, but only for native-like huPrP from *L. tarentolae* including membrane-anchoring. It should be noted that full-length huPrP from *E. coli* is not completely soluble under these conditions so far-UV CD spectra may report on the secondary structure of a mixture of oligomers or aggregates in solution. The higher solubility and a low amount of aggregates and oligomers for native-like huPrP from *L. tarentolae* is supported by SVAUC experiments, revealing the monomer as the most prevalent species (86 % of total signal) as well as a presumably dimeric state. In contrast, full-length huPrP from *E. coli* was detected with only a small number of monomers (21.8 % of total signal) and a large number of aggregates (about 71 % of total signal), which sediment already during acceleration of the centrifuge. From that we conclude that native-like huPrP expressed in *L. tarentolae* is superior to full-length huPrP expressed in *E. coli* with regard to studies under near-physiological conditions. We demonstrated that native-like huPrP expressed in *L. tarentolae* combines the desirable advantages of high protein yields with the presence of PTMs that are close to their human counterparts. A purification protocol was presented, which yields highly pure native-like huPrP solubilized in DDM micelles (Figure 4), where the vast majority is in a monomeric state. Furthermore, these conditions allow for direct transfer into controlled membrane environments such as nanodiscs, as previously shown for amyloid-β precursor protein C99 (Krishnarjuna et al., 2024) and the ion-channel accessory subunit barttin (Viennet et al., 2019). Isotopic labelling of native-like huPrP for NMR studies recombinantly produced in *L. tarentolae* could likely be employed by already established protocols for this organism (Foldynová-Trantírková et al., 2012; Niculae et al., 2006).

**Figure 4.**
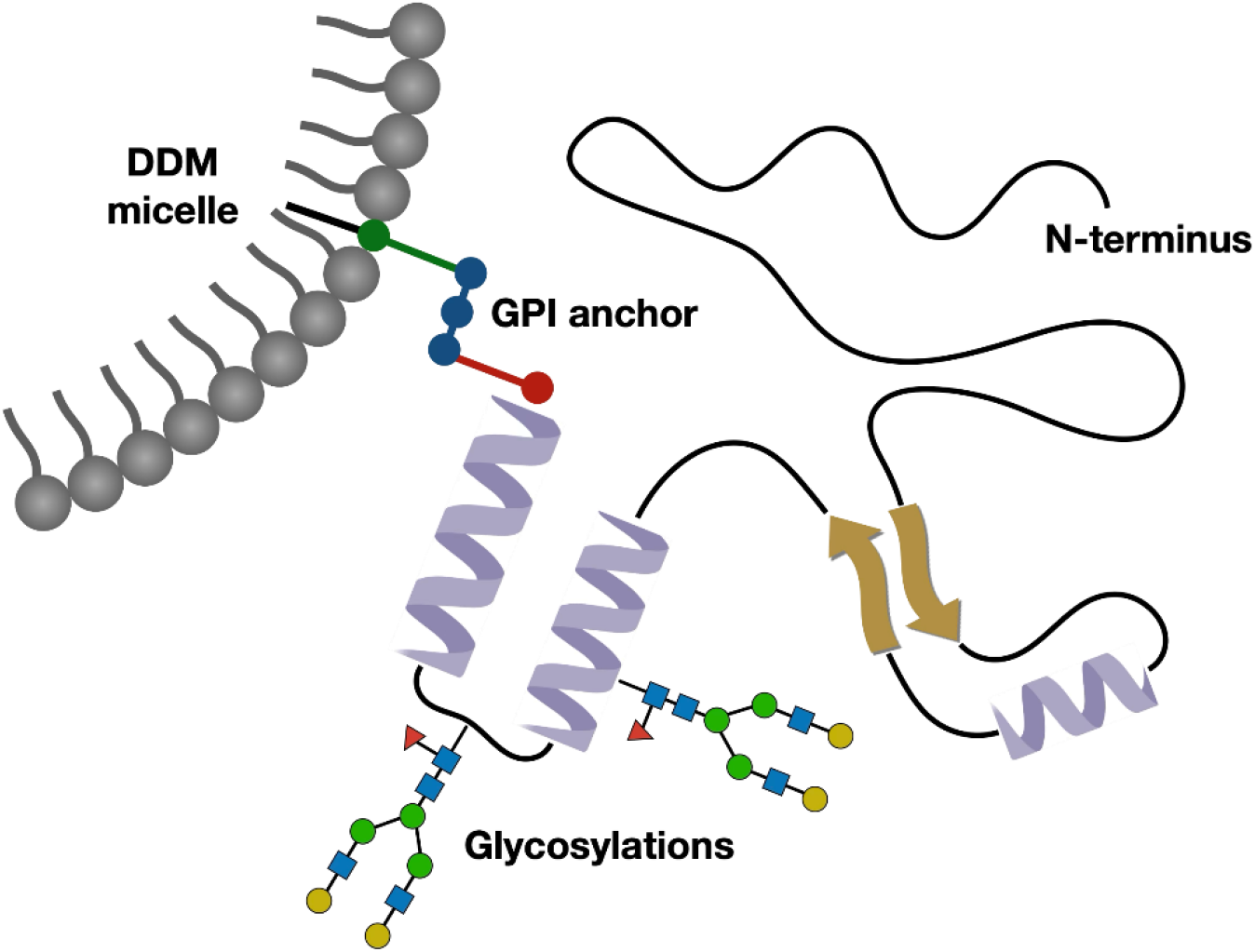
Schematic figure of native-like GPI-anchored huPrP solubilized in DDM micelles. Using the *L. tarentolae* expression system (LEXSY) and the described purification protocol, ∼1 mg/l culture of purified full-length GPI-anchored huPrP is obtained in its natively folded monomeric state, solubilized in DDM micelles.

Thus, elucidating the structural details of native-like huPrP^C^ in its physiologically relevant environment paves the way for future studies of disease-related processes such as its conversion to PrP^Sc^ and mediation of other neurodegenerative diseases through interaction with disease-related proteins.

## Methods

### Cloning of huPrP and amplification of pLEXSY_l-neo2.1

The amino acid sequence was designed as shown in Figure 1A. HuPrP DNA coding sequence (129 Met variant of the naturally occurring polymorphism) was optimized to *L. tarentolae* codon usage (GeneArt—Thermo Fisher Scientific, Regensburg, Germany). Gene synthesis and cloning into the pLEXSY_l-neo2.1 vector, using NcoI/NotI cloning sites, was performed by the company GenScript. NcoI is used for intracellular expression, while NotI was chosen to remove the 6xHis stretch from the backbone. Flanking regions of 5’ CC and 3’ GCGGCCGC were added to form NcoI and NotI sites on the insert. For plasmid amplification, competent XL-1 *E. coli* were transformed with 0.8 µg plasmid. Recombinant *E. coli* clones were selected using ampicillin at 30 °C for plasmid stability reasons. Plasmid identity was confirmed by sequencing (Microsynth, Göttingen, Germany) using 5’-CCGACTGCAACAAGGTGTAG-3’ as forward primer and 3’-CATCTATAGAGAAGTACACGTAAAAG-5’ as reverse primer. After sequence confirmation, 9 µg plasmid was purified from a 50 ml *E. coli* culture using the QIAGEN Plasmid Midi Kit (cat. no. 12143, Qiagen, Hilden, Germany).

### Preparation of the expression plasmid for transfection of L. tarentolae

The huPrP was expressed using the inducible LEXSY expression system according to the manufacturer’s instructions (LEXSinduce Expression Kit, Jena Bioscience, Jena, Germany). 9 µg pLEXSY_l-neo2.1 plasmid including the huPrP gene was digested using SwaI, to remove the 2 kbp *E. coli* fragment. Efficiency was confirmed by agarose gel electrophoresis. Enzymes and buffer salts were subsequently removed using the NucleoSpin Gel and PCR Clean-up Kit (Macherey-Nagel, Düren, Germany). The remaining linearized expression cassette was eluted using 50 µl 10 mM Tris-HCl buffer, pH 8 and used for the transfection of *L. tarentolae*.

### Cultivation and transfection of L. tarentolae

Culturing of *L. tarentolae* was performed according to the manufacturer’s instructions (LEXSinduce Expression Kit, Jena Bioscience, Jena, Germany). Briefly, *L. tarentolae* was cultured in brain heart infusion (BHI) medium (Jena Bioscience, Jena, Germany), supplemented with 5 µg/ml hemin (Jena Bioscience, Jena, Germany), 0.5 % Pen-Strep (Jena Bioscience, Jena, Germany), 100 µg/ml nourseothricin (NTC) (Jena Bioscience, Jena, Germany), and 100 µg/ml hygromycin (Hygro) (Jena Bioscience, Jena, Germany) for the maintenance of T7-TR polymerase and TET repressor genes. Culturing of 10 ml cultures was performed quiescent in 75 cm^2^ ventilated tissue culture flasks at 26 °C in the dark and under aerated conditions. For the cultivation pre-and post-electroporation 10 % fetal calf serum (FCS) (Jena Bioscience, Jena, Germany) was added. After an optical density of 1.3 (approx. 6 × 10^7^ cells) was reached, the cells were centrifuged at 2,000 ×*g* at RT for 3 min and resuspended in half of the remaining medium. Cells, plasmid, and electroporation cuvette with a path length of 2 mm were incubated on ice for 10 min. 350 µl cells were mixed with 50 µl plasmid and incubated in the electroporation cuvette for 10 min on ice. Electroporation was conducted at 450 V and 450 µF for 3.76 msec. After another incubation step on ice for 10 min, the electroporated cells were transferred into a 75 cm^2^ ventilated tissue culture flask, containing 10 ml BHI medium (+ hemin, NTC, Hygro, Pen-Strep, and 10 % FCS). Electroporated cells were incubated at 26 °C in the dark for 6 h before 5 ml of the culture was diluted into 15 ml BHI medium (+ hemin, NTC, Hygro, Pen-Strep, 10 % FCS, and 50 µg/ml selection antibiotic geneticin G-418). Subsequently, the culture was transferred into a 96-well plate, with 200 µl per well. After sealing the plate using Parafilm, it was incubated at 26 °C in the dark. After one week dense cultures became visible and contents of wells with viable cells were transferred into 10 ml ventilated tissue flasks. Genomic DNA integration was tested, by diagnostic PCR, after genomic DNA extraction using phenol/chloroform extraction. Primer pairs for diagnostic PCR were provided by the manufacturer (LEXSinduce Expression Kit, Jena Bioscience, Jena, Germany). Forward primer for the 5’ odc-utr1 control region was 5’-TCCGCCATTCATGGCTGGTG-3’, reverse primer for the 5’ odc-utr1 control region was 5’-TATTCGTTGTCAGATGGCGCAC-3’. Forward primer for the antibiotic-3’ odc control region was 5’-GGATCCAATATGGGATCGGCCATTG-3’ and the reverse primer for the antibiotic-3’ odc control region was 5’-GTGCACCCATAGTAGAGGTGC-3’. This resulted in 1.1 kbp and 2.4 kbp fragments, respectively. After validation of genomic integration, larger volumes (50-600 ml) were cultured in Erlenmeyer flasks at 26 °C in the dark, under aerated conditions, and under agitation at 130 rpm. The T7 promoter-driven transcription was induced by the addition of tetracycline at a final concentration of 15 μg/ml. After 48 h, cells were harvested with centrifugation at 3,000 ×*g* at RT for 10 min.

### Analysis of protein expression

Western blotting was performed to analyze whether the expressed huPrP is localized intracellularly or extracellularly. For intracellular expression analysis, a 2 ml culture with an optical density of 1.0 was centrifuged at 3,000 ×*g* at RT for 10 min. Pelleted cells were resuspended in 0.2 ml reducing SDS-PAGE sample buffer. For secretory protein expression analysis, 8 ml culture was centrifuged at 3,000 ×*g* at RT for 10 min. The supernatant was mixed with 2 ml of 50 % trichloroacetic acid (TCA) and incubated on ice for 30 min. Then, the supernatant was centrifuged at 15,000 ×*g* at 4 °C for 15 min. The pellet was washed with ice-cold 80 % acetone and centrifuged again at 15,000 ×*g* at 4 °C for 15 min. 80 µl of reducing SDS-PAGE sample buffer was added to the pellet. Cell and supernatant samples were incubated at 95 °C for 10 minutes before being applied on SDS-PAGE.

### SDS-PAGE and western blot analysis

For analysis samples were loaded onto a 12 % Tris/Glycine SDS-PAGE at 120 V and subsequently transferred onto a PVDF membrane (PALL Life Sciences, Port Washington, New York) *via* a semi-dry transfer unit (TE70X, Hoefer, Bridgewater, Massachusetts). After blotting, the membrane was blocked for 1 h with 5 % milk powder in Tris-buffered saline containing 0.1 % Tween-20 (TBS-T), washed once with TBS-T, and incubated with anti-PrP antibody (SAF32, 0.1 μg/ml, Bertin Bioreagent, Montigny le Bretonneux, France) in TBS-T overnight at 4 °C. The membrane was then washed three times with TBS-T before secondary antibody goat anti-mouse HRP conjugate (0.1 μg/ml; Jackson Immuno Research Inc., West Grove, Pennsylvania) was added and incubated for 1 h at RT. After three washing steps, the protein detection was performed using a SuperSignal West Pico Chemiluminescent Substrate Kit (Thermo Fisher Scientific, Waltham, Massachusetts). Visualization was conducted on a LAS 4000 (Fuji Film). For dot blot, a concentration series was applied on a dry nitrocellulose membrane. The starting concentration was 0.85 µg of protein in 2 µl buffer (50 mM Tris-HCl, 150 mM NaCl, 0.02 % DDM, pH 7.4), and was further diluted 1:2 in the same buffer. After that, the membrane was blocked with 5 % milk powder in TBS-T and the following steps were identical to the western blot.

### Cell lysis and membrane isolation

After centrifuging of 600 ml *L. tarentolae* culture with an optical density of 1.73, at 3,000 ×*g* at RT for 10 min, the pellet was resuspended in 10 ml lysis buffer (0.33 M sucrose, 0.15 M Tris-HCl, 0.1 M aminocaproic acid, 1 mM EDTA, pH 7.4). Afterwards, glass beads were added into a 15 ml falcon tube, and cells were lysed using a cell disruptor (FastPrep 24; MP Biomedicals, Eschwege, Germany) for 3 times 20 s at 4 m/s. Subsequently, differential centrifugation was conducted to isolate the membrane fraction. First, the cells were centrifuged at 3,000 ×*g* at 4 °C for 5 min, to remove large cell debris and glass beads. Then, the supernatant was centrifuged at 5,000 ×*g* at 4 °C for 10 minutes, followed by an ultracentrifugation step of the supernatant at 100,000 ×*g* at 4 °C for 1 h. The pellet contained the membrane fraction which was used for the solubilization of native-like huPrP.

### Solubilization of membrane-bound native-like huPrP

The pellet containing the membrane fraction was suspended in 6 ml buffer (50 mM Tris, 150 mM NaCl, 1 % DDM, pH 7.4). The sample was incubated for 3 h at 4 °C before it was centrifuged at 100,000 ×*g*, 4 °C for 1 h. Aside from DDM the following conditions were also tested in order to achieve solubilization: denaturing agents and detergents: 6 M Guanidinium chloride (Gdn-HCl), 6 M Urea, 1 % SDS; non-denaturing, non-ionic detergents: 1 % Triton X-100, Tween-20, Igepal; non-denaturing, zwitterionic detergents: 1 % CHAPS, 1 % Zwittergent 3-14; non-denaturing, anionic detergents: 1 % Sodium cholate. The solubilization results with the respective agents and detergents are shown in Suppl. Figure 2. The samples were incubated for 1 h at 4 °C under shaking before they were centrifuged at 100,000 ×*g* at 4 °C for 1 h. Finally, the supernatant (containing the solubilized native-like huPrP) was separated from the pellet (containing non-solubilized huPrP) and both fractions were analyzed by western blot as detailed above.

### GPI-anchoring analysis

For analysis of the GPI-anchoring, 20 µl of supernatant after the 5,000 ×*g* centrifugation step was incubated with 40 µl 0.1 M HEPES-NaOH buffer, pH 7.6, 20 µl of 0.8 % sodium deoxycholate and 20 µl of 1 nM phosphatidylinositol-specific phospholipase C enzyme from *Bacillus cereus* (PI-PLC) (Thermo Fisher Scientific, Waltham, Massachusetts) in 0.1 % BSA for 1 h, 300 rpm at RT. Afterwards, samples were centrifuged at 5,000 ×*g* for 10 min at RT. PI-PLC is supposed to specifically cleave phosphatidylinositol into two molecules, resulting in the release of GPI-anchored proteins from the membrane. Resulting pellets and supernatants were afterwards analyzed by western blotting as detailed above.

### Purification

After solubilization with 1 % DDM, the supernatant was separated by immobilized metal affinity chromatography (IMAC) using a 5 ml Protino Ni-NTA column (Macherey-Nagel, Düren, Germany) after equilibration of the column with 50 mM Tris-HCl, 150 mM NaCl, 0.1 % DDM, pH 7.4. The protein was eluted with a linear gradient from 5 mM to 500 mM imidazole, 50 mM Tris-HCl, 150 mM NaCl, 0.1 % DDM, pH 7.4, within 100 ml. The fractions containing native-like huPrP were pooled and subsequently concentrated using a centrifugal concentrator (Vivaspin 2, Sartorius, Göttigen, Germany) with a 10 kDa cut-off at 2,000 ×*g* at 4 °C. After reaching a total volume of 400 µl, size exclusion chromatography (SEC) was used for further purification. The concentrated fractions were loaded on a Superdex 200 Increase 10/ 300 GL column (Cytiva, Freiburg, Germany) and separated using 50 mM Tris-HCl, 150 mM NaCl, 0.02 % DDM, pH 7.4.

After each step, analytical samples were collected for analysis by SDS-PAGE. Samples for SV-AUC experiments, far-UV CD measurements, and dot blot were taken directly after the SEC from the second peak with elution volume from 14.5 ml to 16.5 ml. Concentration was determined using UV absorbance spectroscopy and a theoretical extinction coefficient of 57995 M^-1^ cm^-1^ at 280 nm. Recombinant huPrP (23-230) from *E. coli*, was expressed and purified as described previously (König et al., 2021; Rösener et al., 2018).

### Far-UV Circular Dichroism (CD) Spectroscopy

Far-UV CD spectra of 10 µM huPrP obtained from *L. tarentolae* or *E. coli* were recorded in a Jasco J-815 (Jasco, Tokyo, Japan) spectropolarimeter. 100 µl sample of the SEC fraction including the native-like huPrP (in 50 mM Tris-HCl, 150 mM NaCl, 0.02 % DDM, pH 7.4) and full-length huPrP from *E. coli* (in 10 mM Tris-HCl, pH 7.4) were filled into a 1 mm quartz glass cuvette and measured as 10 accumulations at 20 °C, 50 nm/min scanning speed, with 2 nm bandwidth and 4 s digital integration time.

### Analytical ultracentrifugation (AUC)

Sedimentation velocity (SV) experiments were performed in an analytical ultracentrifuge Proteome Lab XL-A (Beckman-Coulter, Brea, US). Samples of 10 µM huPrP from *L. tarentolae* or *E. coli* (in 50 mM Tris-HCl, 150 mM NaCl, 0.02 % DDM, pH 7.4) were measured in standard double sector cells (Aluminum) with an optical path length of 12 mm using an An-60Ti rotor. The duration of temperature equilibration to 20°C was 1 h. Data was recorded at 50,000 rpm for 5.5 h. Data analysis was performed using a continuous distribution Lamm equation model, *c*(*s*), implemented in the software Sedfit (version 16p35) (Schuck, Perugini, Gonzales, Hewlett, & Schubert, 2002). Graphical output was created using the software Datagraph (Adalsteinsson, David and Schultz, 2020). In order to eliminate the potential effect of DDM micelles on huPrP solubility regardless of the expression system, huPrP expressed in *E. coli* was additionally measured in the same buffer as native-like huPrP expressed in *L. tarentolae*. No effect of DDM on the solubility of huPrP obtained from *E. coli* was observed.

## Acknowledgment

We gratefully thank Ci Chu, Robin Backer, Linda Reinecke and Christoph Hölbling (all Physikalische Biologie, Heinrich-Heine-Universität Düsseldorf) for technical support.

## Disclosure of conflicting interests

The authors report no conflict of interest.

## Notes

### Competing Interest Statement

The authors have declared no competing interest.

