## Supplemental Figures for "High-Level Expression, Purification and Biophysical Characterization of GPI-anchored native-like human Prion Protein using *Leishmania tarentolae*"

### Supplementary Figures

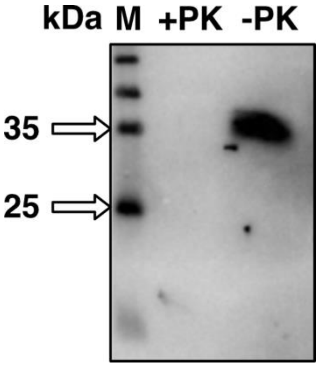


**Supplementary Figure 1. Proteinase K resistance of full-length native-like huPrP from *L. tarentolae*.** Proteinase K digestion of native-like huPrP results in complete digestion. Samples of 50 µl after cell disruption and subsequent 5,000 ×*g* centrifugation were incubated with 50 ng/µl proteinase K for 1 h at 37°C and 300 rpm shaking. Samples were subsequently analyzed by western blotting.

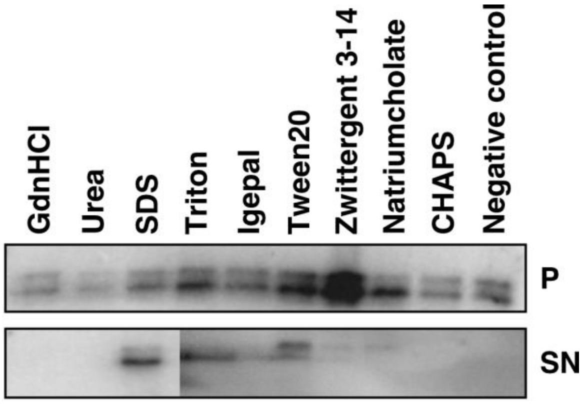


**Supplementary Figure 2. Solubilization of native-like huPrP obtained from from *L. tarentolae* by different denaturants and detergents.** Solubility of native-like huPrP using different detergents. Samples were incubated for 1 h at 4 °C. 6 M GdnHCl and Urea and 1 % of remaining detergents were used for solubilization. “P” represents the pellet and “SN” represents the supernatant after centrifugation for 1 h at 4 °C at 100.000 ×*g*. Samples were analyzed by western blot. Molecular masses were between 25 kDa and 35 kDa.

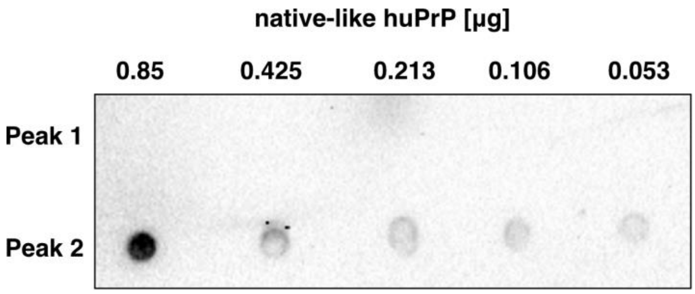


**Supplementary Figure 3. Immunological detection of native-like huPrP from *L. tarentolae* after purification by SEC.** Dot blot of native-like huPrP after SEC. The fractions eluting between 8 ml and 10 ml (Peak 1) as well as 14.5 ml and 16 ml (Peak 2) were concentrated as described for IMAC samples before SEC. A concentration series was applied, starting from 0.85 µg in 2µl and further diluted 1:2 with buffer (50 mM Tris, 150 mM NaCl, 0.02 % DDM, pH 7.4). The lower row shows the second elution peak (Peak 2), verifying the identity of native-like huPrP, while the upper row represents the first elution peak showing no signal for native-like huPrP.

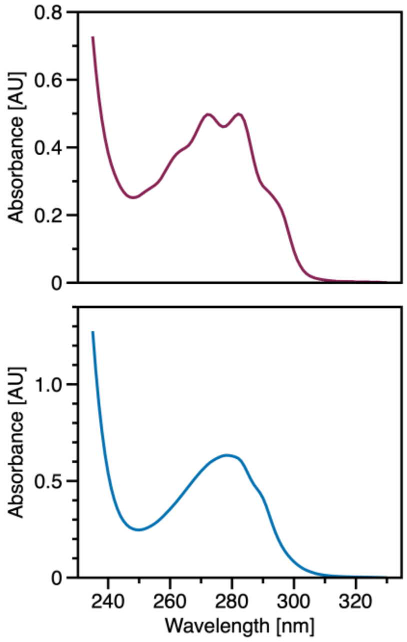


**Supplementary Figure 4. Characteristic UV absorbance spectrum of huPrP obtained from *L. tarentolae* or *E. coli*.** UV absorbance spectroscopy shows distinct absorbance spectra for native-like huPrP expressed in *L. tarentolae* compared to full-length huPrP expressed in *E. coli*. The UV absorbances of ~100 µM purified huPrP obtained from *L. tarentolae* (purple) (in 50 mM Tris-HCl, 150 mM NaCl, 0.02 % DDM, pH 7.4), or from *E. coli* (blue) (in 10 mM Tris-HCl, pH 7.4) are shown.
